# Intrinsic Phenotypic Differences, Not Hemodynamic Shear, Associate with Cusp-Specific Remodeling in Aortic Valve Disease

**DOI:** 10.1101/2025.09.29.679396

**Authors:** Daniel Chaparro, Asad Mirza, David Balzora, Yu Fahong, Valentina Dargam, Lucas Menendez, Ana Valentin, Ian Chen, Sharan Ramaswamy, Joshua Hutcheson

**Author notes:** Dr. Sharan Ramaswamy passed away before the submission of this manuscript. His input and effort were instrumental in this project. May he rest in peace.

## Abstract

Aortic valve disease (AVD) is asymmetric. Various clinical reports indicate that the non-coronary cusp (NCC) is disproportionally burdened by pathological remodeling like inflammation, fibrosis, and calcification. This has long been attributed to the resulting differences in hemodynamic load that arise from the presence, or lack thereof, of coronary ostia in respective sinuses. However, there is little to no empirical evidence to suggest that these differences in hemodynamic shear alone are enough to drive the difference in pathological remodeling that is observed. C57BL/6J mice exhibit a high variance in right coronary ostium (RCO) positioning with respect to the right coronary cusp (RCC). Through computational fluid dynamics (CFD) simulations of a mouse aortic valve (AoV) during end diastolic loading, we show negligible differences in wall shear stress (WSS) between a high RCO on the RCC and no ostium on the NCC. Also, though CFD analyses depict at least an order of magnitude difference in WSS through physiologically relevant ostia positions within the sinuses, ostium position does not correlate with calcification burden in a CKD mouse model of calcific AVD. Cusp dependent extracellular matrix (ECM) abundance analysis reveals asymmetric collagen and elastin content in healthy adult mice, but this does not follow the same trend as pathological remodeling. Instead, asymmetric abundance of elastin (P=0.034) and collagen (P=0.018) was mainly driven by an increase of these ECM proteins in the left coronary cusp (LCC) with no differences in leaflet thickness. Cusp dependent transcriptomic (spatial and bulk RNA sequencing) analyses reveal asymmetric phenotypic profiles between the three cusps in healthy adult mice. Of note, genes associated with vascular smooth muscle cell contraction and known modulators of AoV remodeling were upregulated in the NCC and downregulated in both the RCC and LCC. Together, these data suggest that the differences in shear resulting from the coronary ostia are not sufficient to explain the asymmetric onset of calcific AVD.

## Introduction

The aortic valve (AoV) sits between the left ventricle and the aorta, promoting unidirectional blood flow throughout the cardiac cycle. It is normally composed of 3 cusps that open during systole and close during diastole by coapting radially to close the aortic orifice, preventing regurgitant flow into the left ventricle. This happens roughly 3 billion times in a lifetime necessitating proper biomechanical compliance and an ability to withstand tensile, bending, and hemodynamic shear loads [1]. Aside from hemodynamic shear on the aortic side of the cusps during diastolic loading, all other loading conditions are generally thought to be radially symmetric [2]. The diastolic asymmetry is due to the presence of the coronary ostia. During diastole, as blood rushes back from the aorta and closes the AoV, some of the blood flows into the coronary arteries by way of the coronary ostia, which are proximally superior to the cusps. However, there are only two ostia. One behind the aptly named left-coronary cusp (LCC) and one behind the right-coronary cusp (RCC) while the non-coronary cusp (NCC) lacks an ostium. Therefore, while the bending and tensile loads may be similar, the hemodynamic environment imparted on the aortic surface of the cusps, because of the ostia, differs between the LCC, RCC, and NCC [1]. Increased flow into the ostia during diastole increases the wall shear stress (WSS) the surrounding tissues experience. While a lack of an ostium results in lower WSS on the tissue due to reduced and disturbed flow patterns [1, 3].

A hallmark of the pathological progression of aortic valve disease (AVD) is calcification. Along with other pathological remodeling like inflammation and fibrosis, calcific nodule formation on the AoV cusps impairs valve function, leading to stenosis and/or regurgitation, which may necessitate surgical intervention to alleviate cardiac burden and prevent heart failure [4]. Clinical observations consistently report an asymmetric onset of pathological remodeling between the NCC, LCC, and RCC of the AoV during AVD [5]. The NCC is on average more calcified than the LCC and the RCC [3, 6–9]. In the vasculature, atherosclerotic plaque and calcific nodule formation are often found in areas of disturbed, low shear flow such as bifurcations and bends in the vascular tree [10]. In the context of the AoV, pathological blood velocities and pressures can impact bulk tissue properties and influence pathological differentiation and remodeling [11]. It is therefore rational to conclude that these differences in pathological remodeling seen across the AoV cusps must be a direct cause of the presence of the coronary ostia. However, a direct comparison between ostia presence, position, the resulting wall shear stress on the cusps, and cusp calcification burden has not yet been assessed.

In this study, through computational fluid dynamics (CFD) simulations of the mouse AoV during end diastolic loading, we show evidence indicating that the resulting differences in WSS between a high right coronary ostium (RCO) on the RCC and no ostium on the NCC are negligible. Also, though CFD analysis depicts an order of magnitude difference in WSS across physiologically relevant ostia positions within the sinuses, ostium position does not correlate with calcification burden in a chronic kidney disease (CKD) mouse model of AVD. Since CFD predicted WSS loads on the leaflet surface were unable to predict or follow similar trends to observed pathological calcification, we aimed to determine if baseline levels of ECM protein abundance or intrinsic transcriptomic profile differences between the three cusps would give a more informed insight into the asymmetric onset of disease progression prior to initiation.

## Methods

### Animal studies

In this study, 4-week-old male (n = 54) and female (n = 53) mice on C57BL/6J background were used. All mice were bred and housed on a 12:12-h day/night cycle at a room temperature (20 ± 2°C) in the Florida International University Animal Care Facility. The mice were given *ad libitum* access to food and water. All experimental procedures were approved by the Institutional Animal Care and Use Committee at Florida International University under protocol 24-006 and conformed to the Guide for the Care and Use of Laboratory Animals [National Institutes of Health (NIH), Bethesda, MD, United States] for scientific purposes.

Mice were divided into a diseased group (n = 61) and a control group (n = 46). The control group was fed a normal chow diet (5V75 - PicoLab® Verified 75 IF, LabDiet) for 12 weeks and serve as a negative control. The diseased group received the same standard show diet supplemented with high-adenine (0.2%) diet for 6 weeks to induce CKD, followed by a high-adenine, high-phosphorous (1.8%) diet (CKD+HP) for an additional 6 weeks to induce AoV calcification. We have previously shown that the CKD+HP diet induces both medial vascular and valvular calcification in male and female mice [12–14].

### Aortic Valve Dissections

Mice were euthanized by CO2 asphyxiation followed by cervical dislocation. Hearts were resected and kept in ice cold 1 X phosphate buffered saline (1XPBS) until further dissection. Hearts were dissected using a Leica S9i stereoscope to gain access to the AoV, while continually bathing the tissue in ice cold 1XPBS to mitigate degradation, as previously described [15]. In short, all anatomical structures except the ascending aorta and left ventricle are removed including the apical portion of the ventricles. The anterior portion of the left ventricle and the mitral valve were removed. An axial cut was made by inserting one blade of fine dissection scissors underneath the anterior leaflet of the mitral valve and sliding it upwards through the ascending aorta. A cut was carefully made along the LCC and NCC commissure to expose the AoV cusps (**Error!** Reference source not found.).

### Aortic Valve Anatomical Measurements

We took images of the intact AoV apparatus once it was fully dissected, and the cusps were exposed using the Leica S9i stereoscope and LAS X Life Science Microscope Software Platform. These images were then imported into ImageJ to annotate anatomical landmarks (i.e., points). The points were carefully placed at the intersect between each cusp annulus and the virtual basal ring at the LCC-RCC and RCC-NCC interleaflet triangles, at the commissures, at both ends of the aortic root just below the STJ, and at the center of the ostia (**Figure 1**). The coordinates of these points were imported into MATLAB, where a custom script was used to calculate cusp width, STJ height, AoV diameter, and ostia position within the coronary sinuses. Ostium positions were normalized to respective sinus geometry, width and height (**Figure 1 Error! Reference source not found.A**). Cusp width was determined by commissure-to- commissure length, and height was determined by the STJ to annulus length **Figure 1**.

**Figure 1.**
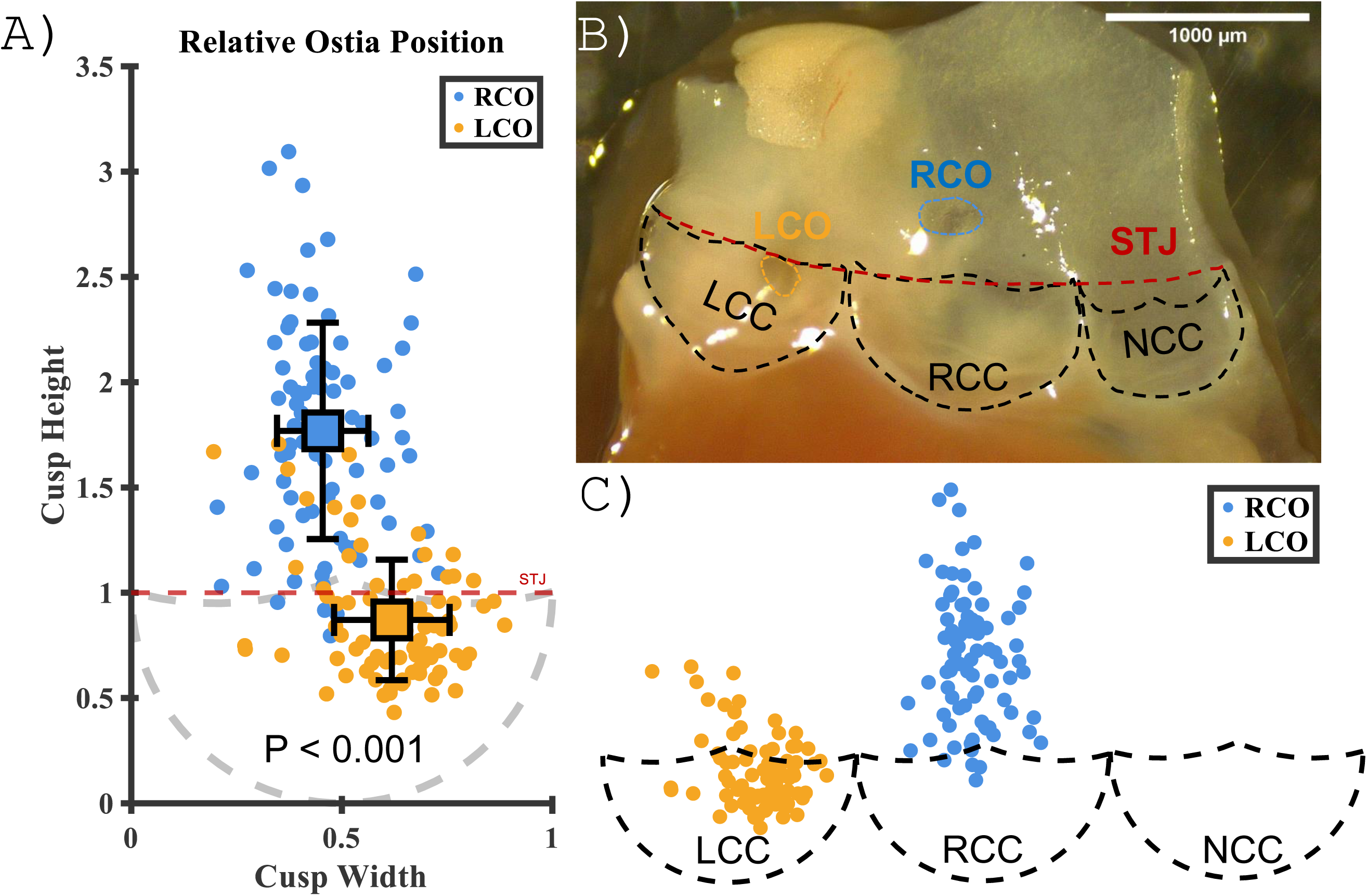

### Computational Fluid Dynamic Simulations

A parameterized, computer aided design (CAD), model of an AoV was created in SolidWorks 2020 (Dassault Systèmes, Vélizy-Villacoublay, France) that included anatomical features such as the annulus, sinus, individual leaflet cusps, coronary ostia, and STJ (**Figure 2**).It was based off similar models developed for analysis of aortic valve disease states [16–18]. While initially created for a human valve, parameters were designed to be universally applicable fit any tricuspid valve in another mammal.

**Figure 2.**
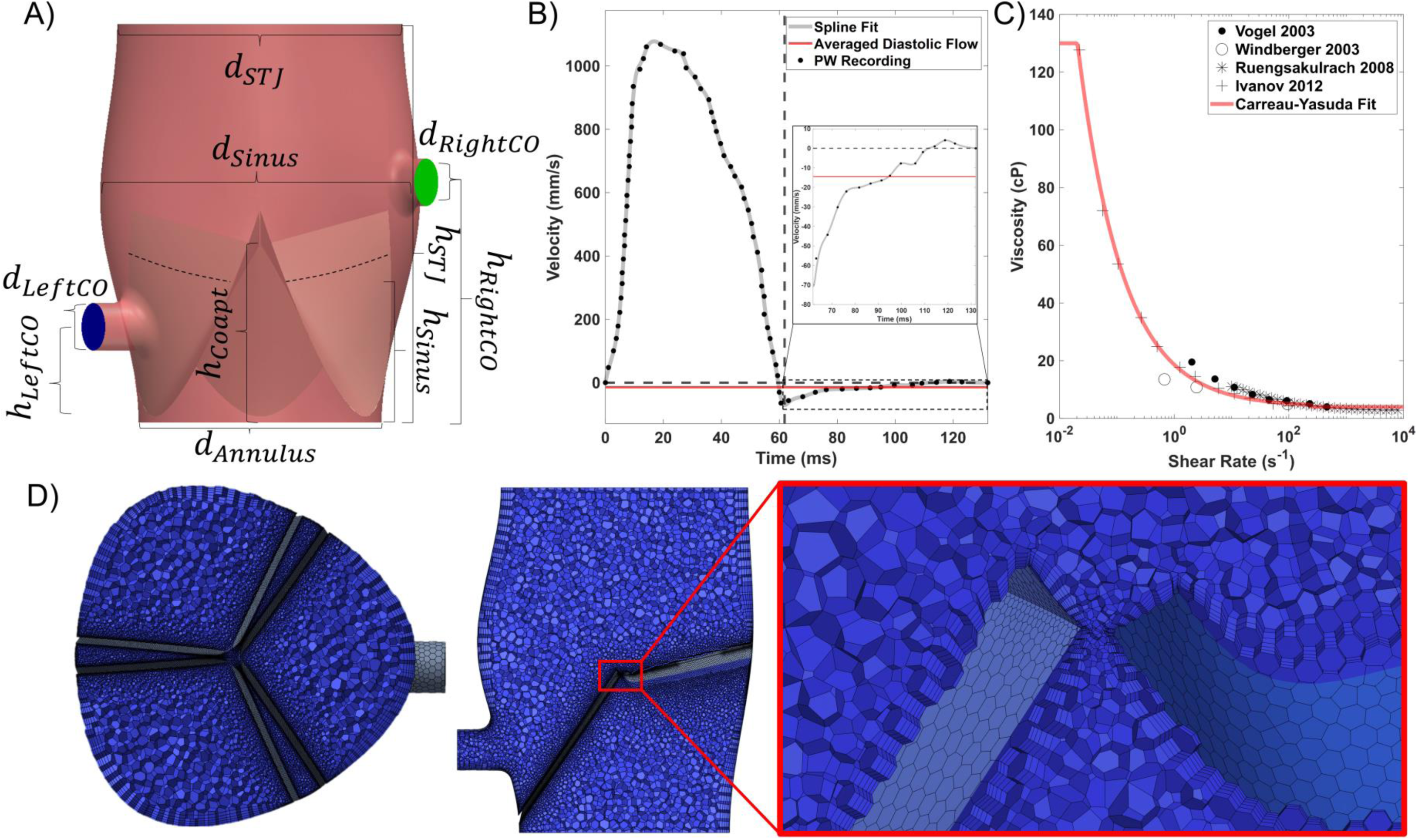

Therefore, mouse AoV anatomical measures acquired were used to calculate averaged dimensions of the major parameters used in the CAD model. Control geometry parameters were as follows with all units being in mm: 𝑑_𝐴𝑛𝑛𝑢𝑙𝑢𝑠_ = 0.868, 𝑑_𝑆𝑖𝑛𝑢𝑠_ = 1.056, 𝑑_𝑆𝑇𝐽_ = 1.02, 𝑑_𝐿𝑒𝑓𝑡/𝑅𝑖𝑔ℎ𝑡 𝐶𝑂_ = 0.17, ℎ_𝑆𝑖𝑛𝑢𝑠_ = 1.423, ℎ_𝐶𝑜𝑎𝑝𝑡_ = 0.631, ℎ_𝐿𝑒𝑓𝑡𝐶𝑂_ = 0.34, ℎ_𝑅𝑖𝑔ℎ𝑡𝐶𝑂_ = 0.859..Right coronary ostium height (ℎ_𝑅𝑖𝑔ℎ𝑡𝐶𝑂_) was varied between 0.3130 and 1.5305 mm. Thus, a total of 6 values of ℎ_𝑅𝑖𝑔ℎ𝑡𝐶𝑂_ were simulated: 0.3130, 0.5565, 0.8590, 1.0435, 1.2870, and 1.5305 mm. Left coronary ostium height (ℎ_𝐿𝑒𝑓𝑡𝐶𝑂_) was also investigated at 6 values: 0.2085, 0.31, 0.417, 0.51, 0.834, and 1 mm. Tri-leaflet symmetry was assumed to reduce computational complexity. Ventricular and aortic inlet and outlets as well as coronary outflow tracts were extended a length 𝑑_𝑆𝑖𝑛𝑢𝑠_ to mitigate any computationally induced entrance or exit effects. Leaflet thickness was assumed to be 45 μm [15]. Prior to meshing, all geometries were exported to ANSYS Spaceclaim 2022 R2 (ANSYS Inc., Canonsburg, PA) for any necessary surface repair that manifested from the exporting process such as closing small gaps, removing small faces, fixing short or inexact edges, and correcting almost tangent faces.

The domain was meshed with polyhedral elements to minimize total number of elements, improve convergence, and reduce solution time (**Figure 2D**). In addition, this element type has been reported to have more homogenous wall metrics than conventional tetrahedral elements [19]. A mesh independence study was conducted on the control geometry parameters to identify the optimal mesh size. Three meshes: coarse, medium, and fine, were created with increasing number of elements each (Table 1). Maximum velocity magnitude in the domain and the surface averaged shear rate (ASR) (Eq. 1) on the belly region of the right coronary leaflet were tracked between meshes.

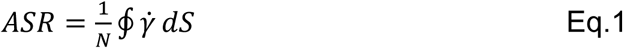

**Table 1:** Mesh independence study

| *Mesh* | Maximum Cell Length (mm) | Number of Elements | Max Velocity During Diastole |  | Average Shear Rate (ASR) |  |
| --- | --- | --- | --- | --- | --- | --- |
| | | | $\vert V_{Max}\vert $ (mm/s) | Percent Diff (%) | ASR (1/s) | Percent Diff (%) |
| 1. Coarse | 0.1000 | 65,736 | 0.81 | 17.45 | 288.55 | 23.90 |
| 2. Medium | 0.0400 | 406,571 | 0.70 | 2.90 | 358.32 | 2.34 |
| 3. Fine | 0.0150 | 831,398 | 0.68 | - | 366.82 | - |

The study resulted in mesh 2, medium size, having less than 5% of a difference in maximum velocity and ASR when compared with the fine mesh, thus a target mesh size of 406,571 elements or a maximum cell length of 0.04 mm was chosen for all further analysis. Cell quality was maintained with an average skewness of 0.08 and an average aspect ratio less than 5 across all meshes. A total of five boundary layers were grown on all wall surfaces to attain more accurate wall metrics. A minimum of three elements was maintained between all close edges, primarily leaflet free edges, excluding boundary layers, to provide adequate room for flow development through the coaptation point (**Figure 2D**), regardless of the maximum cell length imposed on the domain. All geometries were meshed using ANSYS Fluent Meshing.

Two main hemodynamic parameters were tracked in this study: shear rate and wall shear stress, as they have been well established metrics in investigations of calcification in both aortic valves [20–22] and arteries [23, 24]. Shear rate, 𝛾͘, is defined as the gradient, ∇, of the velocity vector, 𝑣⃗ (Eq. 1). Wall shear stress incorporates shear rate and blood’s non-Newtonian viscosity, 𝜇(𝛾͘), (Eq. 2).

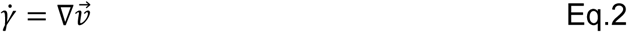

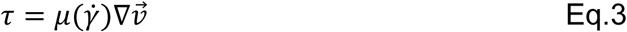

Since wall metrics across the leaflets can vary from tip to base [25] each leaflet was split into 2 regions, the belly and the tip, with the split line located at 10% of the radial length of the leaflet, measured from the base near the annulus (**Figure 2A**).

The steady and incompressible form of the Navier-Stokes equations for continuity and momentum were used to model blood flow (Eq. 3 and Eq. 4):

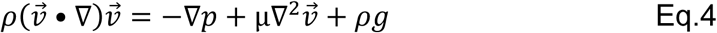

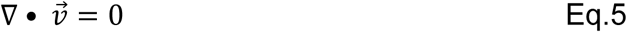

where ∇ is the gradient operator, 𝑣⃗ is the velocity vector, 𝑝 is pressure, 𝜌 is fluid density, 𝜇 is dynamic viscosity, and 𝑔 is the external acceleration due to gravity. Residuals for the continuity and velocity equations were set to 10^-6^. Monitors for ASR and surface averaged wall shear stress (AWSS) (Eq. 6) of belly regions of leaflets were also monitored for convergence.

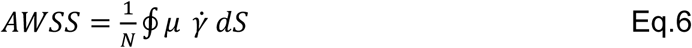

A coupled schema was used for the pressure-velocity solver along with a second-order discretization for the pressure and momentum. Relaxation factors of 0.5 were enforced for the pressure and momentum equations to improve solution stability. Conservative Reynolds number calculations using given information about aortic annulus diameter, blood density, viscosity, and expected peak velocity during diastole indicates peak values of ∼5. This value is well below the threshold for turbulent flow and therefore a laminar model was used for the solver. A similar calculation for the Womersley number was done to ascertain the degree of oscillatory flow and showed a value of ∼ 0.75, indicating that laminar, Poiseuille-like flow, should be expected. The finite volume method in ANSYS Fluent 2022 R2 was used for all simulations.

A single, biphasic, cardiac cycle, from pulsed wave doppler recordings, of ventricular inlet velocity was digitized and segmented using MATLAB R2023b (Mathworks, Natick, MA). Specifically, the diastolic region of the cycle was extracted and fitted with a smoothing spline (**Figure 2B**, gray), this new continuous velocity waveform was then averaged to a constant value of -14.55 mm/s (**Figure 2B**, red) and prescribed on the inlet. Aortic outflow was set to an average downstream aortic pressure of 84.2 mmHg from measurements of wildtype mice [26]. For the left and right coronaries, there were no pressure or flow recordings available from mice so average coronary pressures during diastole from a human [27] were used instead with left and right pressures of 84.23 and 85.46 mmHg, respectively. All other surfaces were treated as walls with a no-slip condition.

While murine and human blood are both shear thinning fluids their viscous behavior differs slightly due to variations in cellular components such as neutrophils and lymphocytes [28]. As such, rather than using a more readily available human model, a murine specific Carreau-Yasuda model (Eq. 7) was fitted from available data [29–32] using a non-linear least-squares algorithm in MATLAB R2023b. This model defines flow between two regimes at various shear rates (𝛾͘): where 𝜇_0_ and 𝜇_∞_ are for viscosity at zero and infinite shear rates, respectively, alongside empirically determined values of λ, α, and *n*. Fitted parameters were 𝜇_∞_ = 3.995 cP, 𝜇_0_ = 130 cP, 𝜆 = 48.82 s, 𝑎 = 75.47, and 𝑛 = 0.4465. Blood density was set to a constant 1057 kg/m^3^ [33].

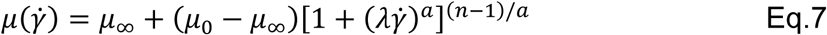

Simulations were performed on a dedicated workstation for simulations with a 4 core, 2661 MHz, 2.67 GHz Intel® Xeon® X5550 (Intel Corporation, Santa Clara, CA) with a NVIDIA Quadro FX 1800 (Nvidia Corporation, Santa Clara, CA) with 16 GB of RAM. CAD modeling was done in SolidWorks 2020 with cleaning and mesh repair in ANSYS SpaceClaim 2022 R2 under Windows 10 Enterprise 64-bit (Microsoft Inc., Redmond, WA) Postprocessing and data analysis was done in ANSYS EnSight 2022 R2 and MATLAB R2023b.

We used these results to generate predictive models for AWSS on the aortic surface of each cusp during end diastolic loading given the position of the coronary ostia. A predictive model for each cusp was independently determined using a linear interpolation method (**Figure 3B**) with MATLAB built in function *fit*. These models were then used to estimate the AWSS on each cusp of all the mice in the study with relevant AoV anatomical measurements (**Figure 3C**).

**Figure 3.**
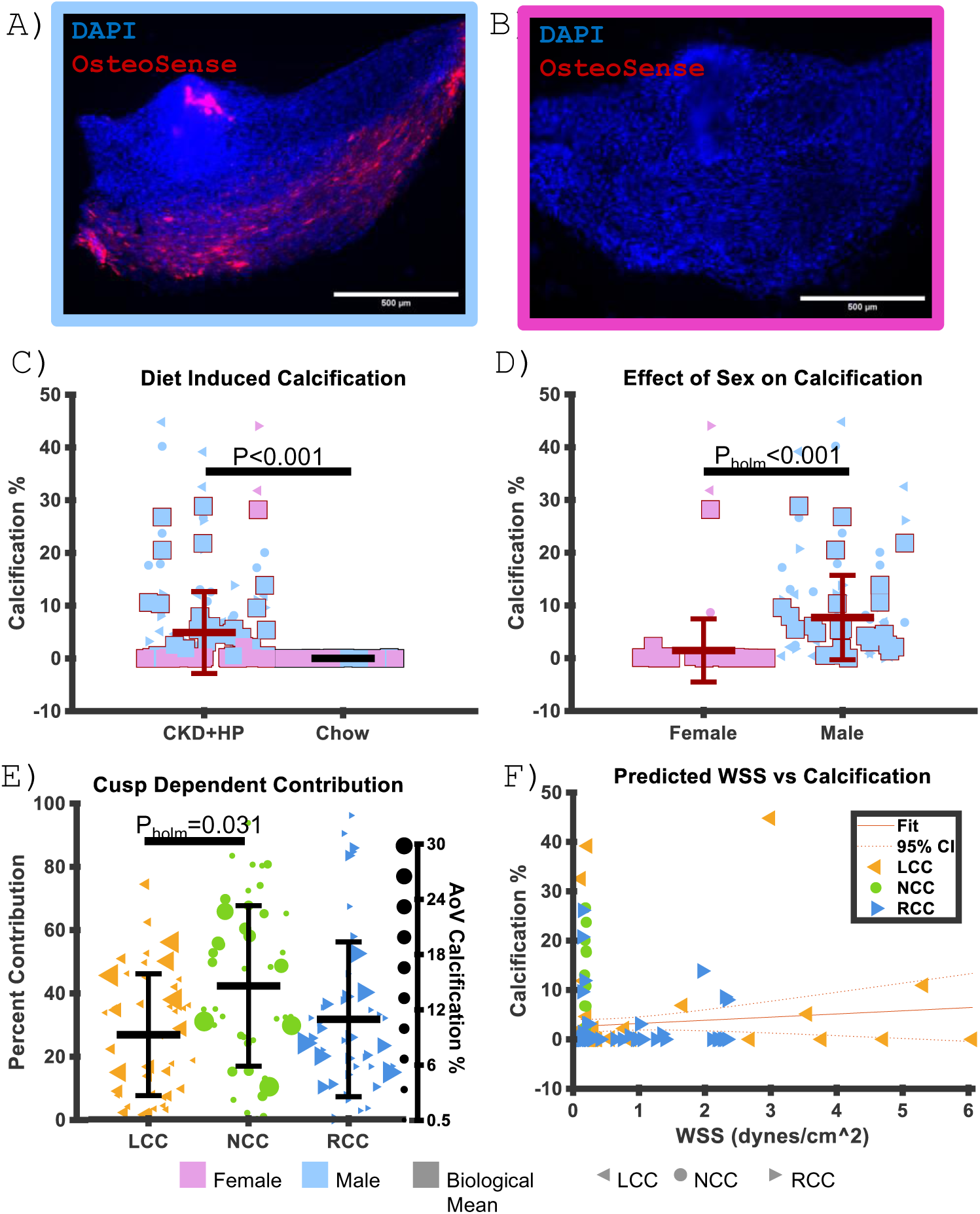

### Calcification Quantification

After 12 weeks on their respective dietary regimens, 16-week-old mice were injected via tail-vein with OsteoSense680x (80 nmol/kg, Perkin Elmer, Waltham, MA, United States) one day prior to euthanasia to visualize calcification deposits on valvular tissues.

OsteoSense is a calcium mineral binding probe widely used to image calcification in the cardiovascular field [12, 34]. Of the 107 mice in the study, 72 mice were used for this purpose. AoVs from 49 mice on the CKD+HP diet (27 males and 22 females) and 23 mice from the chow diet (10 males and 13 females) were resected as previously described. One male and one female from the CKD+HP group were excluded from analysis due to bicuspid AoV malformations. Once the cusps were exposed, the tissue was fixed in 4% paraformaldehyde (PFA) at room temperature for 10 minutes and then placed in 1xPBS in the fridge (4°C) overnight. The following day, cusps were resected and mounted on microscope slides. We used a Zeiss Axioscope epi-fluorescent scope equipped with filter sets for DAPI (Filter Set 49 Zeiss) and OsteoSense680x (49022 ET – Cy5.5 Chroma). A custom MATLAB script was used to quantify the area of OsteoSense positive signal based on a manually chosen threshold and normalized to the area of the tissue calculated by a binarized image from the DAPI signal. AoV calcification was quantified as the fraction positive OsteoSense signal. For cusp dependent analyses, only calcified valves were included. Only valves which calcified more than 3 standard deviations away from the chow fed mean were included. All the valves included were exclusively from the CKD+HP group.

To assess how coronary ostia position within the sinuses and other relevant AoV anatomical parameters would predict cusp calcification, we ran linear regression models using the MATLAB function *regress* between the percentage of cusp calcification and ostia height (LCC and RCC only), cusp width, cusp height, and AoV diameter. These analyses were performed on leaflets from valves which calcified at least more than 3 standard deviations from the chow fed mean.

### Extracellular Matrix Abundance Quantification

To understand other potential underlying differences between the cusps and to assess the effect of the CKD+HP diet on ECM remodeling, we stained mouse AoV cusps with elastin and collagen probes. The remaining 35 mice were used for this purpose. AoVs from 12 mice on the CKD+HP diet (6 males and 6 females) and 23 mice on the chow diet (11 males and 12 females) were resected as described above. One male from the CKD+HP group was excluded from analysis due to a bicuspid aortic valve malformation. Once the cusps were exposed, the tissue was fixed in 4% PFA in 1xPBS for 10 minutes at room temperature and then placed in 1xPBS in the fridge (4°C) overnight. We stained the tissues the following day with DAPI, a fluorescent collagen probe (CNA35-OG488) and an elastin probe (Alexa Fluor 633 hydrazide) to image nuclei and structural ECM fibers in AoV tissues [35, 36]. DAPI, collagen and elastin probes were diluted in 1xPBS (1µg/mL, 1µM and 0.2µM respectively), placed on the tissues, and incubated at room temperature for 30 minutes. The tissues were subsequently rinsed 5 times with 1xPBS. We resected out each individual cusp and mounted them on microscope slides for imaging. A Nikon C2 confocal microscope with a 60x oil immersion objective was used to acquire optical sections (512x512 pixels at 0.41µm/pixel) of the belly region of each cusp through the entire thickness of the tissue. We wrote a custom MATLAB script to quantify abundance of each fluorophore per AoV cusp as the mean fluorescence intensity (MFI) and normalized to the volume of the tissue imaged. Relative cusp contributions to thickness, elastin, and collagen content were calculated by normalizing the values from each cusp to its respective biological mean value. All statistical analyses were done with repeat measures ANOVA of the non-normalized values in JASP.

### Spatial and Bulk RNA Sequencing

Aside from gross anatomical and hemodynamic loading differences, we wanted to understand if there were underlying phenotypic differences between the cusps that could exacerbate pathological disease progression prior to the onset of CKD or calcification. Two 4-week-old mice (1 male and 1 female) were used for spatial transcriptomic analysis of AoV cusp tissue as previously described [37]. A transverse cut was made to remove the apex of the heart. Atria, pulmonary artery, and right ventricle were carefully resected. Finally, the remaining posterior aspect of the left ventricle and mitral valve were removed. The remaining tissue (interventricular septum, anterior aspect of left ventricle and mitral valve, AoV, and aortic root) were snap frozen in a liquid nitrogen cooled bath of isopentane. The tissues were then embedded in 7×5×5 mm disposable base molds in OCT ensuring that the long axis of the tissue was perpendicular to one of the mold faces and then stored at -80°C.

Tissue molds were placed in a cryostat at -20 °C for sectioning. Molds were sectioned proximal to the tissue to prevent OCT overlap within a capture area of the 10X Visium Spatial slides (10X Genomics, Pleasanton, CA). Serial caudal transverse sections of the tissue were made starting from the superior aspect of the aortic root and placed on microscope slides to determine when the AoV was reached. Once at the AoV, serial 16 μm sections of tissue were placed on the 10X Genomics Visium Spatial slides. Two to six consecutive sections were placed on a single capture area. Four capture areas were used for each biological sample. We followed 10x Genomics “Methanol Fixation, H&E Staining and Imaging - Visium Spatial Protocols” protocol (CG000160) to fix and stain the tissues on the slides. A Zeiss Axio Observer.Z1 microscope was used to create large, high resolution (0.91 μm/pixel), stitched images of the capture areas ensuring the fiducial frame of each capture area was clearly visible.

For the library construction, 25% of the total cDNA samples obtained from the capture areas were used. cDNA was eluted with buffer EB, followed by fragmentation and adaptor ligation. Post ligation cleanup using SPRIselect (Beckman Coulter Cat# B23317), PCR-based sample indexing, and quality analysis using Agilent bioanalyzer high sensitivity chip were performed. Illumina NovaSeq6000 platform was used with 25 million reads per area. Sequencing was done on S4 flow cell lanes using 300 cycles.

We used 10X Genomics Loupe Browser visualization software to manually select the AoV leaflet tissues within the tissue sections and distinguish between LCC, NCC, and RCC tissue (298 unique spots). The corresponding barcodes to the manually selected spots corresponding to the AoV leaflet tissue were imported into MATLAB for subsequent analysis. A custom MATLAB script was used to align the raw read count data set with the spatially identified AoV leaflet barcoded spots. Only genes that had at least 10 read counts were included (∼9,000 genes). MATLAB function *rnaseqde* was used to determine differentially expressed genes (DEGs) between comparisons of interest. T- distributed stochastic neighbor embedding (t-SNE) dimensionality reduction analysis was done with the MATLAB function *tsne* with default settings and random number generator set to *“default”* for reproducibility.

AoV cusps from an additional 7 chow fed 4-week-old mice (4 females and 3 males) were resected for cusp specific bulk RNA sequencing. After each dissection, each cusp was carefully removed from the AoV apparatus, placed in 350 µL of lysis buffer with 5µL of 4ng/µL carrier RNA, and homogenized using a sonicator following the manufacturers recommendation (QIAGEN RNeasy micro-Kit). The samples were placed on ice and transferred to a -80°C freezer. The following day, samples were thawed on ice and proceeded with the RNA purification protocol outlined by the manufacturer (QIAGEN RNeasy Micro Handbook March 2021). The 10µL eluates were placed in a Styrofoam shipping container with dry ice and shipped to the University of Florida’s Interdisciplinary Center for Biotechnology Research (UF | ICBR) for library construction and bulk RNA sequencing.

Reads acquired from the Illumina NovaSeq 6000 platform were cleaned up with the cutadapt program (Martin 2011) to trim off sequencing adaptors and low-quality bases with a quality phred-like score <20. Reads <40 bases were excluded from RNA-seq analysis. The genome of Mus musculus (version GRC38, mm10) from the Ensembl database was used as the reference sequences for RNA-seq analysis. The cleaned reads of each sample were mapped to the reference sequences using the read mapper of the STAR package (Spliced Transcripts Alignment to a Reference, v2.7.9a) [38].The mapping results were processed with the HTSeq (High-Throughput Sequence Analysis in Python, v0.11.2) [39] samtools, and scripts developed in house at ICBR of UF to remove potential PCR duplicates and count uniquely mapped reads for gene expression analysis. PCA analysis (for detecting outlier samples) and volcano plot analysis based on all identified genes in each analysis were performed with the R-package (v4.1.3).

Differential expression between groups of interest was done with the MATLAB function *rnaseqde*. For leaflet dependent comparisons, each leaflet was treated as a biological replicate. For example, of the 21 leaflets, when comparing the NCC to the RCC and LCC, the 7 NCCs were assigned as the treated group and the 14 other leaflets were assigned as the control group. Genes with an FDR adjusted p-value of less than 0.05 and a 1.5x fold change were considered significantly differentially up/down regulated genes in that leaflet compared to the other two. Lists of DEGs were then loaded into Cytoscape for network analysis [40]. Networks were constructed using the StringApp protein query application [41]. StringApp was also used to generate functional enrichment tables with GO Biological Process, STRING Clusters, and KEGG Pathways as references. The lists of pathways were then imported into MATLAB to determine intersecting pathways across the leaflets.

### Computational deconvolution of leaflet bulk RNA-seq

We complemented standard bulk RNA-seq analyses with computational deconvolution to resolve leaflet-specific cell type composition and lineage programs. Bulk cusp samples (7 LCC, 7 RCC, 7 NCC) were harmonized with a postnatal murine aortic valve single-cell reference (Hulin et al., GEO: GSE117011) [42], which includes endothelial, interstitial, immune, and melanocyte lineages. Gene names were standardized to official symbols, version suffixes were removed, and low-count features were excluded.

Duplicates were collapsed by total count, and both bulk and single-cell matrices were intersected to a shared gene universe to ensure consistency of features.

Single-cell reference data were filtered, normalized, log-transformed, and clustered using canonical valve markers (e.g., Pecam1, Cdh5 for endothelium; Col1a1, Dcn for interstitial; Tyrobp, Ctss for immune; Tyr, Pmel for melanocyte). Pseudo-bulk reconstructions of P7 and P30 stages reproduced expected developmental differences, confirming suitability of the reference for deconvolution.

Leaflet deconvolution was performed with Tissue-AdaPtive autoencoder (TAPE), a deep autoencoder framework for tissue-adaptive mixture analysis [43]. TAPE models bulk expression as a linear combination of non-negative cell type signatures, with an encoder to estimate fractions and a decoder that outputs a gene-by-cell-type signature matrix. Training was performed in two stages: (1) simulated mixtures generated from the single-cell reference, including both typical and sparse compositions, with inputs scaled to [0,1] per gene; and (2) per-sample adaptation in high-resolution mode to capture subtle leaflet context. Final outputs included estimated cell type fractions and a sample-specific signature matrix representing predicted lineage expression programs.

Leaflet asymmetry was assessed using TAPE fractions. Because TAPE outputs are compositional, fractions were transformed using a centered log-ratio (CLR) transformation per sample:

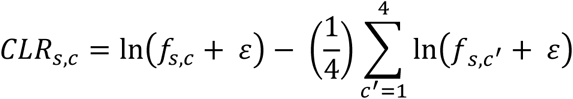

Leaflet differential abundance (Δ) was defined for each cell type *c* and leaflet L ∈ (LCC, RCC, NCC) as the mean in L minus the mean of the other two leaflets.

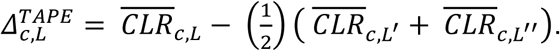

Per-sample contrasts were compared using Welch’s t-test with Benjamini–Hochberg adjustment for 12 contrasts (4 cell types × 3 cusps). To account for inter-mouse variability, per-mouse Δ values were also computed:

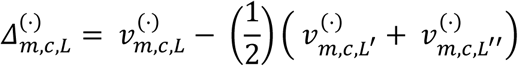

with *v^TAPE^ = CLR*.

## Results

### Ostia Position

LCO and RCO positions within respective sinuses are significantly different (P<0.001, Wilcoxon signed rank test of paired distances between ostia). The LCO tends to originate at the level of the sinotubular junction and near the LCC-RCC commissure while the RCO tends to originate above the sinotubular junction and slightly off center towards the LCC-RCC commissure (**Figure 1C**). The frequency of high-take-off, defined as an ostium originating above the STJ [44], of the LCO and RCO was 22.4% (13/58) and 72.4% (42/58) respectively. When the LCO was high (N=13) the RCO was also high in 92.3% of occurrences (12/13).

### CFD Simulations

A total of six different RCO and LCO heights were modeled to explore their relation to wall shear stress on each leaflet during diastole. The CFD analyses were all performed under identical initial conditions, boundary conditions, and other physiological parameters. RCO height varied from 0.3130 mm to a maximum of 1.5305 mm with respect to the annulus and LCO height from 0.2085 mm to 1 mm (**Figure 3A** 3D models subset). Surface contours of AWSS on the belly regions of all leaflet cusps are shown for all right coronary heights simulated (**Figure 3A** heatmaps). When RCO and LCO heights were nearly identical, 0.2085 mm for left coronary and 0.3130 mm for right, the velocity vectors and corresponding AWSS on the left and right cusps were also similar regardless of height. When left and right ostium heights were the lowest, the left cusp had an AWSS of ∼17.18 dynes/cm^2^ and right ∼18.87 dynes/cm^2^, respectively. However, as the RCO height increased there is a noticeable drop in AWSS on the RCC, with the final height at 1.5305 mm having an AWSS of ∼0.12 dynes/cm^2^ (**Figure 3B** right graph). This resulted in the wall metrics of the RCC and NCC becoming nearly identical (**Figure 3A** heatmaps). Wall metric distributions for all cusps appear to be independent and only a function of their respective ostium’s height. Most of the drop AWSS occurred between the RCO heights of 0.3130 mm and 0.5565 mm, or just before passing the STJ at 0.631 mm.

Based on our simulations and predictive models of AWSS based on ostia positions, there is cusp dependent asymmetric shear on the aortic surface of the cusps during end diastolic loading (P<0.001 Repeat Measures ANOVA with post hoc holm correction) (**Figure 3C**). On average, the LCC experiences more AWSS than the NCC and RCC (P_holm_<0.001 and P_holm_=0.005 respectively). While the NCC AWSS is lower than the RCC (P_holm_=0.001).

### Diet Induced Calcification

Compared to chow fed controls, the CKD+HP diet significantly increased the amount of detected calcification based on OsteoSense positive signal on the tissue (P<0.001, Two-way ANOVA with post hoc Holm correction) (**Figure 4C**). There was a sex dependent difference in calcification. Males calcified significantly more than females (P_holm_=<0.001) (**Figure 4D**). Cusp contribution to calcification is significantly different (P=0.044, Repeat Measures ANOVA with post hoc Holm correction) (**Figure 4E**). The NCC contributes significantly more to calcification burden than the LCC (P_holm_=0.031), but there is no significant difference within other combinations of comparisons. We also note a bimodal distribution of NCC contribution to calcification. A similar yet less pronounced bimodal distribution can be observed in LCC contribution to calcification as well.

**Figure 4.**
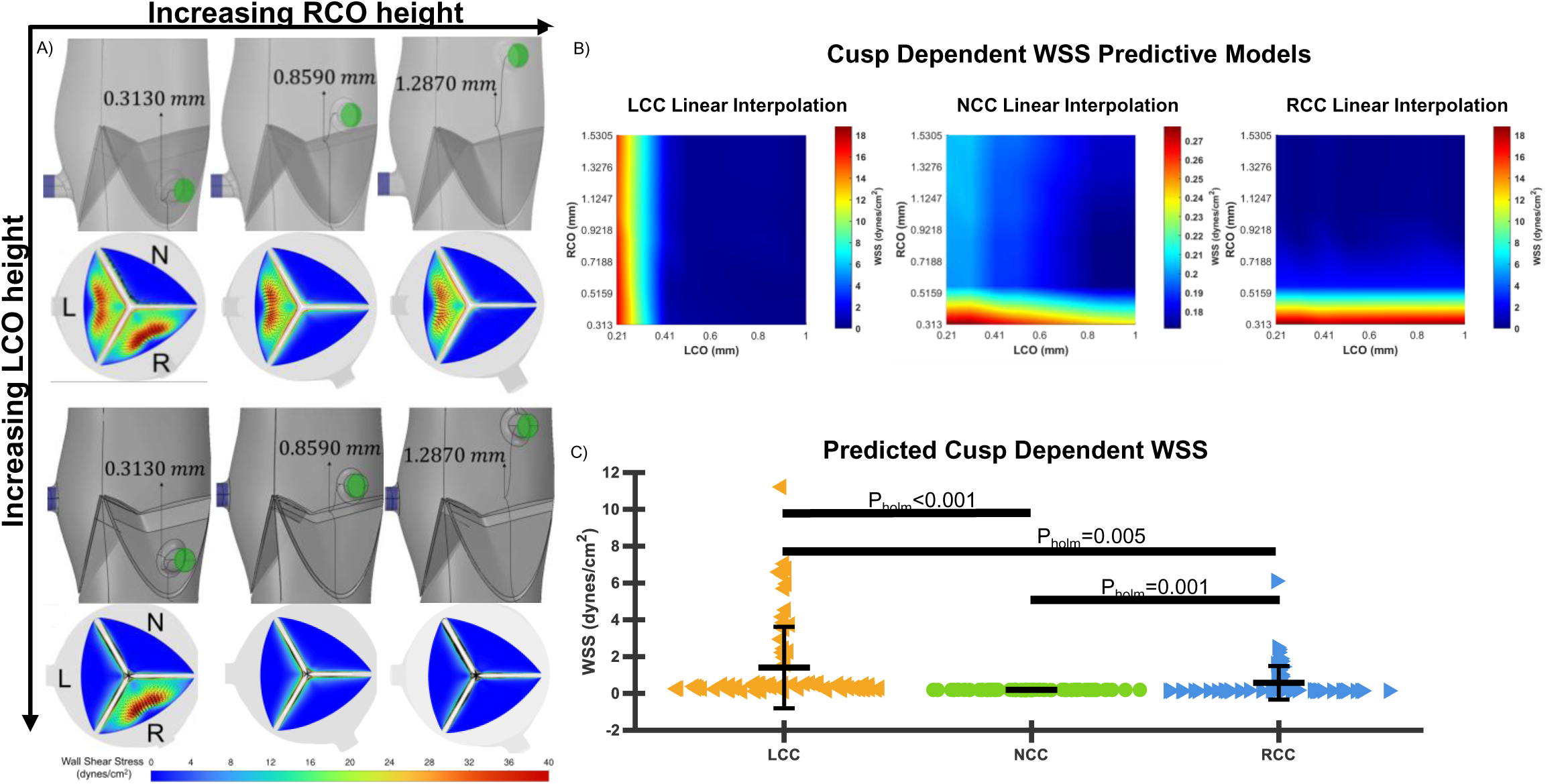

Predicted AWSS did not correlate to calcification burden in either the LCC, NCC, or the RCC (R-Squared=0.002, P=0.793) (**Figure 4F**). Multiple regression analysis of all relevant anatomical features (AoV diameter, cusp width, cusp height) did not result in a statistically significant model to predict calcification (Model R-Squared=0.2033, P=0.451).

### ECM structural Imaging

We were interested in determining baseline differences in ECM composition that could account for the observed calcification response described above. Though female chow fed controls trended to have less elastin abundance than their male counterparts (P=0.071, Welch T-Test), there was no statistically significant sex dependent differences in thickness, elastin content, or collagen. There were, however, cusp dependent differences in elastin (P=0.034) and collagen (P=0.018) content (Repeat Measures ANOVA with Post Hoc Holm correction) (**Figure 5** **C** and **D** respectively). The LCC had more elastin (P_holm_=0.029) and collagen than the RCC (P_holm_=0.041) and tended to have more elastin (P_holm_=0.109) and collagen (P_holm_=0.06) than the NCC, but this did not reach statistical significance (**Figure 5C** and **D**). Interestingly, the RCC and NCC were the most similar in the context of ECM composition. There were no differences or appreciable trends in thickness between the cusps (**Figure 5B**).

**Figure 5.**
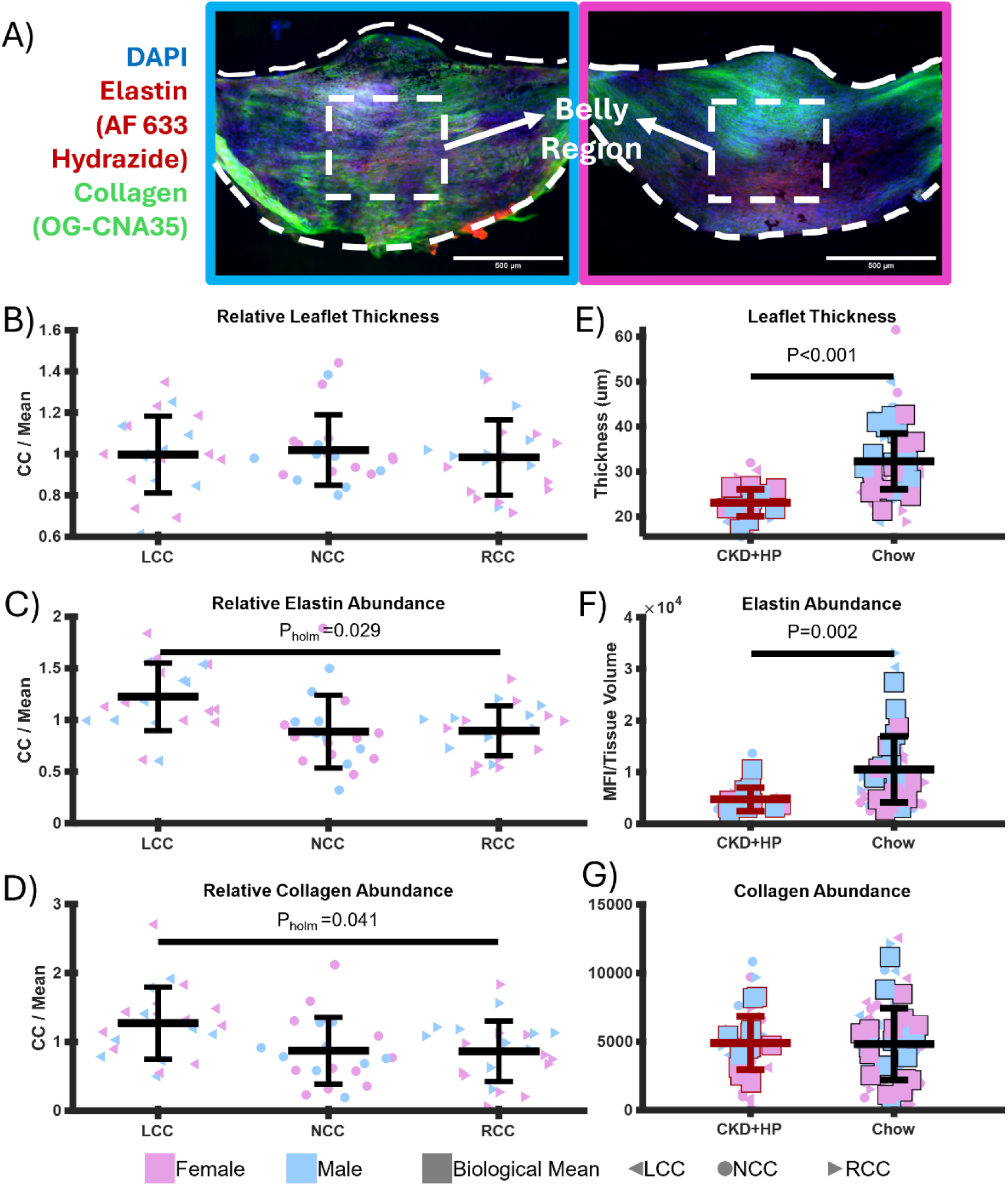

In the context of AVD, compared to chow fed controls, AoV cusps form mice on the CKD+HP diet were significantly thinner (P<0.001, Two-way ANOVA) (**Figure 5E**), and had less elastin abundance (P<0.01, Two-way ANOVA) (**Figure 5F**). Collagen content between the groups was not different (**Figure 5G**). We did not observe any sex dependent differences in these responses.

### Cusp Specific Transcriptomic Profiles

From spatial transcriptomic analyses of two adult valves (1 male 1 female), t-SNE dimensionality reduction depicts that the cusp tissue clusters together when compared to surrounding tissue however, the cusps do not cluster separately from one another (**Figure 6**).

**Figure 6.**
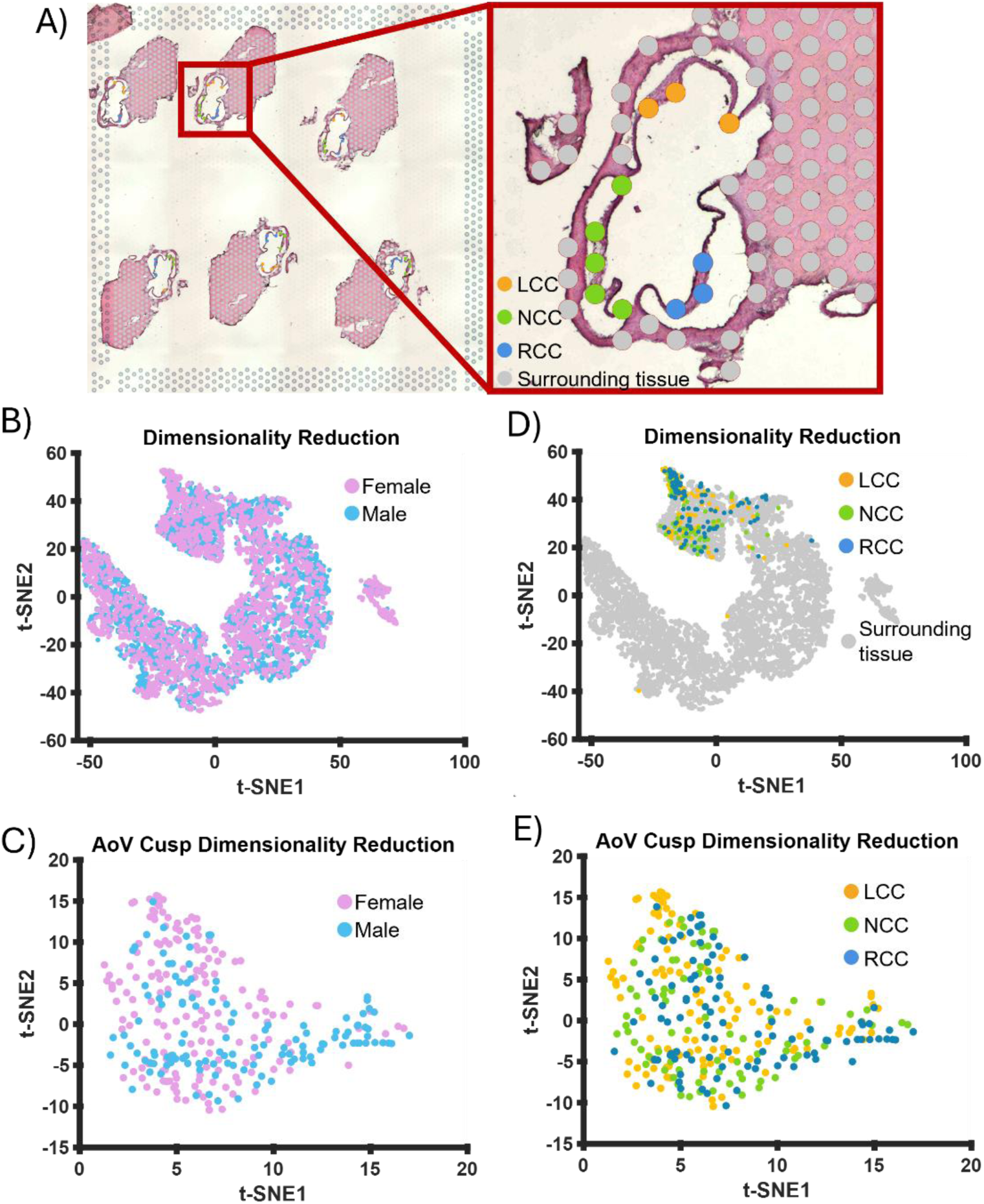

There was no sex-dependent clustering. Comparing transcriptomic profiles of the AoV cusps to that of surrounding tissues yielded 4818 DEGs (FDR Adjusted P-Value<0.05) of which 1129 were at least 2-fold increase in the cusps and 2211 were 2-fold decreased in the cusps. The top 10 up and down regulated genes from this comparison can be found in Table 2. Sex-specific comparison of AoV cusps tissue yielded 1349 DEGs (FDR Adjusted P-Value<0.05) of which 1319 were at least 2-fold increase in the female AoV cusps and 11 were at least 2-fold decreased in the female AoV cusps. The top 10 up and down regulated genes from this comparison can be found in Table 3.

**Table 2:** AoV Cusps vs Surrounding Tissues

| Table 2: AoV Cusps vs Surrounding Tissues |  |  |  |  |
| --- | --- | --- | --- | --- |
| Upregulated in AoV |  |  | Downregulated in AoV |  |
| Gene | Adjusted P-Value |  | Gene | Adjusted P-Value |
| **Postn** | 1.52E-292 |  | **Myl2** | 0 |
| **Wif1** | 1.42E-285 |  | **Mb** | 0 |
| **Coch** | 7.73E-208 |  | **mt-Rnr2** | 0 |
| **Thbs1** | 5.63E-204 |  | **mt-Nd1** | 0 |
| **Vcan** | 2.79E-197 |  | **mt-Nd2** | 0 |
| **Vim** | 4.92E-184 |  | **mt-Nd4** | 0 |
| **Ptn** | 1.94E-164 |  | **mt-Cytb** | 0 |
| **Dkk3** | 1.54E-160 |  | **Myl3** | 5.57E-295 |
| **Ahsg** | 1.05E-154 |  | **Myh6** | 6.96E-256 |
| **Scara3** | 4.81E-138 |  | **mt-Co1** | 6.78E-236 |

**Table 3:** Female vs **Male** AoV Cusps

| Table 3: **Female** vs **Male** AoV Cusps |  |  |  |  |
| --- | --- | --- | --- | --- |
| Upregulated in Female AoV |  |  | Downregulated in Female AoV |  |
| Gene | Adjusted P-Value |  | Gene | Adjusted P-Value |
| **Xist** | 5.84E-09 |  | **Hba-a1** | 1.09E-08 |
| **Ccn2** | 1.80E-05 |  | **Ddx3y** | 7.28E-07 |
| **Gm10076** | 2.45E-05 |  | **Hbb-bs** | 1.88E-06 |
| **Myoc** | 0.000126188 |  | **Hbb-bt** | 0.0028398 |
| **H2-Ab1** | 0.000128416 |  | **Cox8b** | 0.010246862 |
| **Apoe** | 0.000128416 |  | **Myh11** | 0.02195913 |
| **Gsn** | 0.000128416 |  | **Myoz2** | 0.026291224 |
| **Nppa** | 0.000128416 |  | **Mmp3** | 0.028139462 |
| **H2-Aa** | 0.000243013 |  | **Hba-a2** | 0.040914238 |
| **Vim** | 0.000333704 |  | **Retnla** | 0.041261765 |

Comparing the LCC to the other two cusps yielded 1764 DEGs (FDR Adjusted P-Value<0.05) of which 1432 were at least 2-fold increase in the LCC and 6 were at least 2-fold decreased in the LCC. The top 10 up and down regulated genes from this comparison can be found in Table 4. Comparing the NCC to the other two cusps yielded 5 DEGs (FDR Adjusted P-Value<0.05) of which 1 was at least a 2-fold increase in NCC and 4 were at least 2-fold decreased in the NCC. The top 10 up and down regulated genes from this comparison can be found in Table 5. Comparing the RCC to the other two cusps yielded 60 DEGs (FDR Adjusted P-Value<0.05) of which 1 was at least 2-fold increase in RCC and 56 were at least 2-fold decreased in the RCC. The top 10 up and down regulated genes from this comparison can be found in Table 6.

**Table 4:**
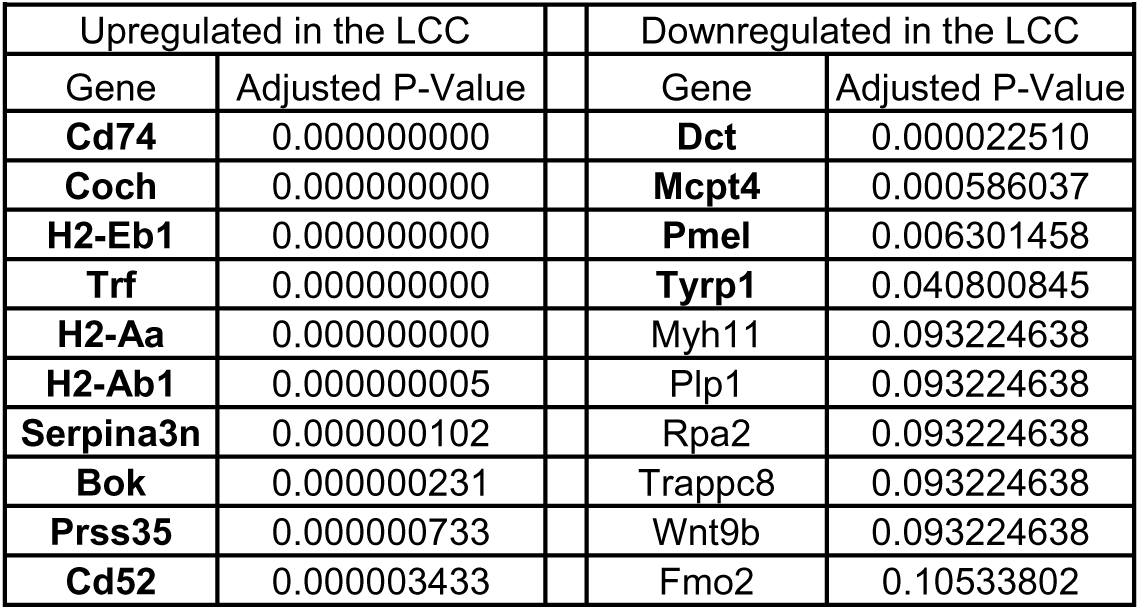
LCC vs **RCC** and **NCC**

**Table 5:** NCC vs **RCC** and **LCC**

| Table 5: **NCC** vs **RCC** and **LCC** |  |  |  |  |
| --- | --- | --- | --- | --- |
| Upregulated in the NCC |  |  | Downregulated in the NCC |  |
| Gene | Adjusted P-Value |  | Gene | Adjusted P-Value |
| **Hba-a1** | 0.006703860 |  | **Myoc** | 0.00152109 |
| Adamts8 | 0.146649348 |  | **Stab1** | 0.013859103 |
| Plp1 | 0.165130662 |  | **Cd74** | 0.036406757 |
| Palmd | 0.357226305 |  | **Bok** | 0.043493392 |
| Myh11 | 0.357226305 |  | Cycs | 0.068449306 |
| Bst2 | 0.357226305 |  | Mb | 0.068449306 |
| Rnf138 | 0.432534344 |  | Trf | 0.068449306 |
| Impdh1 | 0.456725423 |  | H2-Ab1 | 0.068449306 |
| Csnk1g1 | 0.459934921 |  | H2-Aa | 0.068449306 |
| Nf1 | 0.459934921 |  | Coch | 0.084109821 |

**Table 6:** RCC vs **LCC** and **NCC**

| Table 6: **RCC** vs **LCC** and **NCC** |  |  |  |  |
| --- | --- | --- | --- | --- |
| Upregulated in the RCC |  |  | Downregulated in the RCC |  |
| Gene | Adjusted P-Value |  | Gene | Adjusted P-Value |
| **Mcpt4** | 0.000178304 |  | **H2-Eb1** | 1.17E-05 |
| Cma1 | 0.100242054 |  | **Cd74** | 1.17E-05 |
| Dct | 0.102763577 |  | **Coch** | 0.000138554 |
| Zfp518b | 0.216994213 |  | **Trf** | 0.000260006 |
| Rpa2 | 0.229434484 |  | **H2-Aa** | 0.000260006 |
| Pign | 0.259491027 |  | **Bst2** | 0.001230316 |
| Car8 | 0.266947604 |  | **Hbb-bs** | 0.001268462 |
| Smad1 | 0.274241923 |  | **Serpina3n** | 0.002481399 |
| Smcr8 | 0.276409735 |  | **H2-Ab1** | 0.002481399 |
| Tyrp1 | 0.285590534 |  | **Hba-a1** | 0.003200868 |

Cusp specific bulk RNA sequencing analyses of an additional 7 (3 male, 4 female) healthy adult (30 days old) mice revealed more underlying phenotypic differences amongst the cusps that were generally consistent with the findings from the spatial transcriptomic analysis. This is all summarized in **Figure 7**. Comparing the LCCs to the other two cusps yielded 209 DEGs (FDR Adjusted P-Value<0.05) of which 36 were at least 1.5-fold increased in the LCC and 138 were at least 1.5-fold decreased in the LCC (**Figure 7A** left panel). Comparing the NCCs to the other two cusps yielded 310 DEGs (FDR Adjusted P-Value<0.05) of which 223 were at least 1.5-fold increased in the NCC and 55 were at least 1.5-fold decreased in the NCC (**Figure 7A** left panel). Comparing the RCCs to the other two cusps yielded 98 DEGs (FDR Adjusted P-Value<0.05) of which 40 were at least 1.5-fold increased in the RCC and 52 were at least 1.5-fold decreased in the RCC (**Figure 7A** left panel). Network analyses from these sets of DEGs (**Figure 7B and C**) together with functional enrichment analyses resulted in 5 different pathways which were common in at least two distinct cusp DEG clusters: vascular smooth muscle contraction (mmu04270), ossification (GO:0001503), melanogenesis (CL:12121), angiotensin regulation (CL:36185), and response to IFN-β (CL:26940) (**Figure 7D**). Of these, only “vascular smooth muscle contraction” was present in all three of the cusps, appearing in the upregulated NCC cluster and downregulated LCC and RCC clusters.

**Figure 7.**
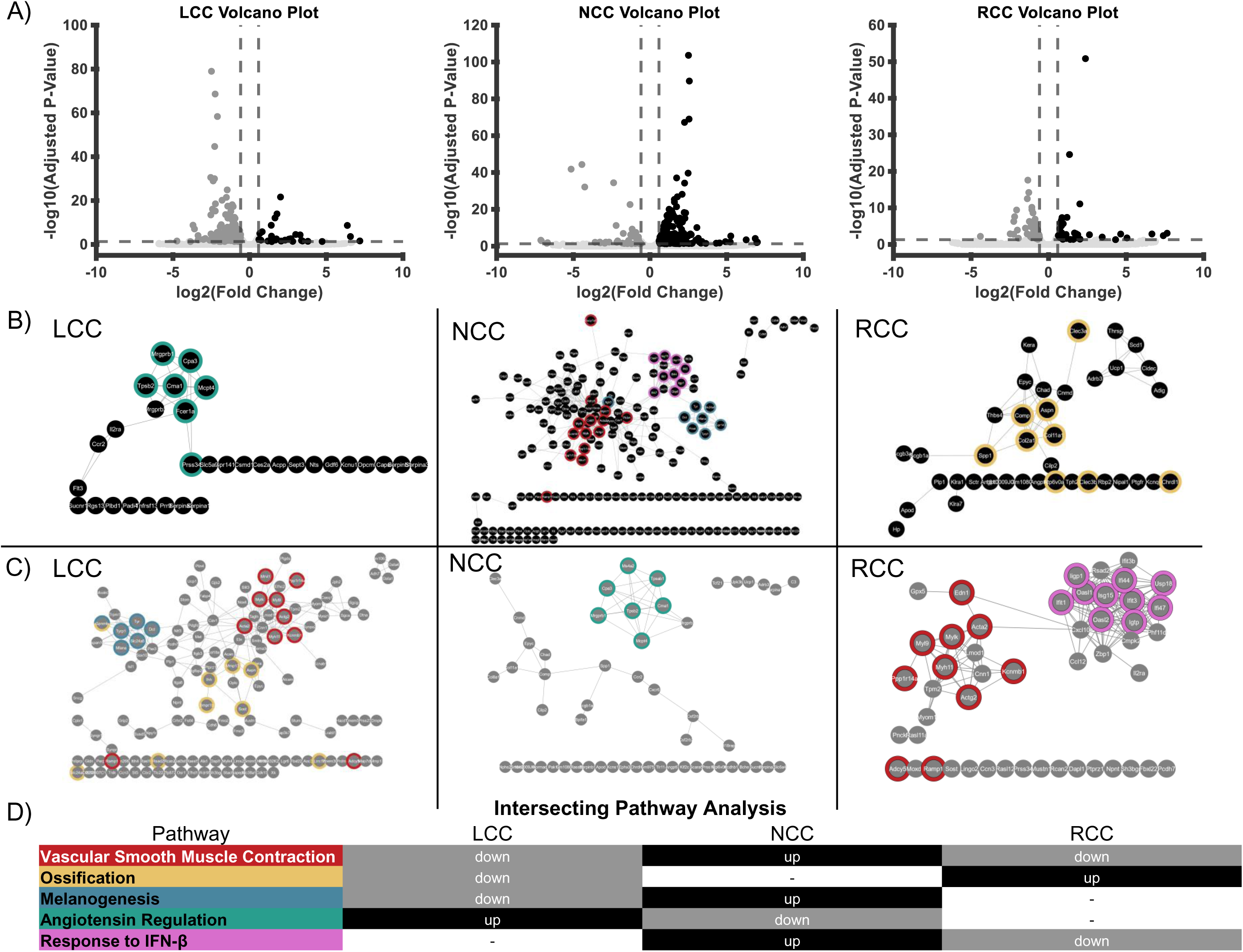

### Computational Deconvolution

Computational deconvolution revealed clear cusp-dependent variation in lineage composition and transcriptional programs (**Figure 8**). After a Greenhouse-Geisser sphericity correction, repeat measures ANOVA showed that there is a statistically significant difference proportions of interstitial cells (P=0.001), endothelial cells (P=0.007), immune cells (P=0.026), and melanocytes being the most significant (P<0.001) between the 3 leaflets. A summary of post-hoc Holm pairwise comparisons for each cell type can be seen in **Figure 8A**. Endothelial and interstitial cells were the dominant lineages across all cusps, but the LCC showed enrichment for interstitial signatures, while the NCC displayed comparatively higher endothelial proportions.

**Figure 8.**
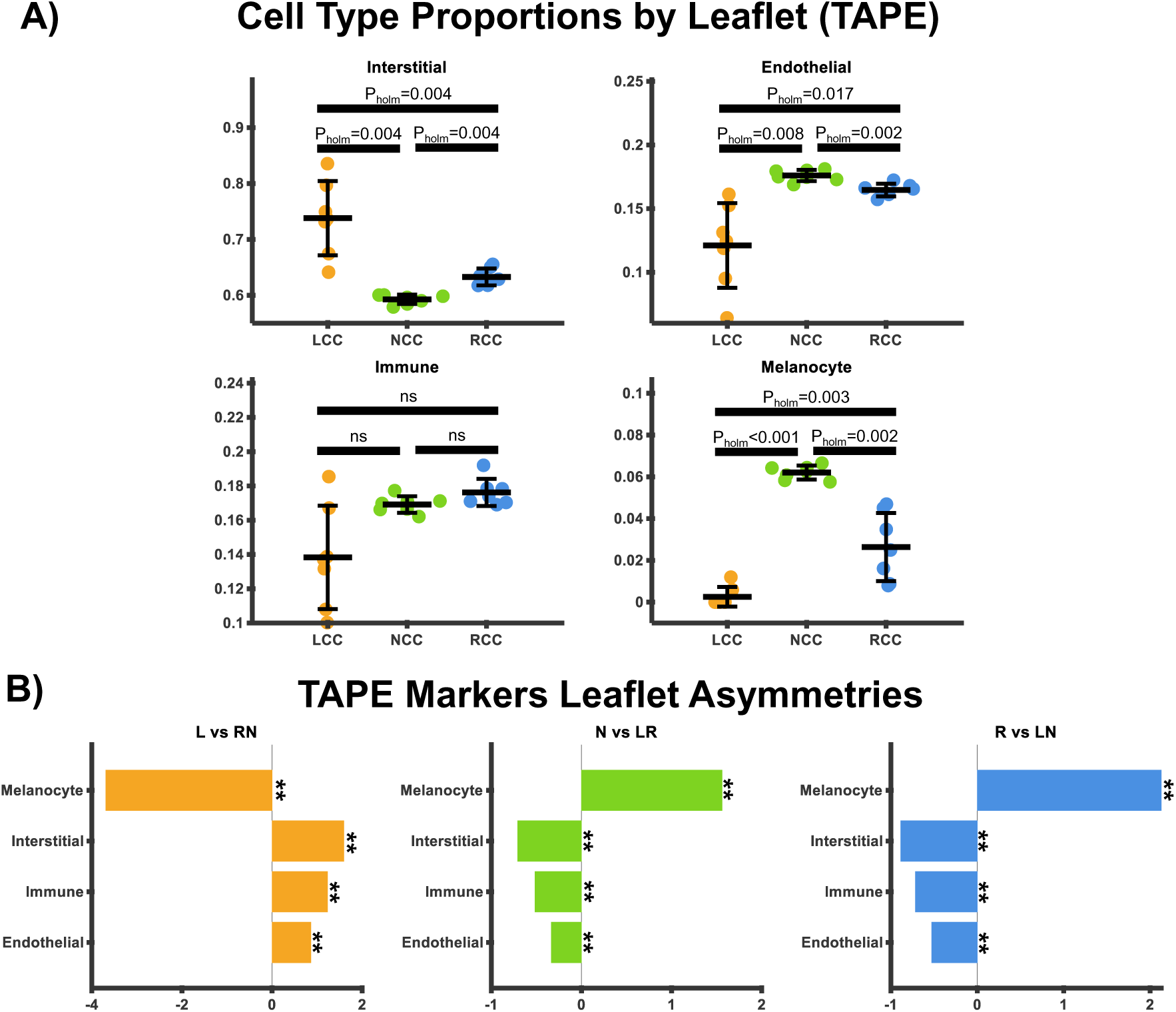

Immune-derived contributions were consistently elevated in the RCC relative to the LCC and NCC but the post hoc Holm pairwise comparisons did not reach statistical significance. The NCC presented with significantly higher melanocytic proportions compared to the LCC and RCC. Leaflet asymmetries, as measured by CLR, depict statistically significant differences for each cell type and each leaflet compared to the other two (e.g. LCC vs NCC + RCC) (FDR adjusted p-value<0.01) (**Figure 8B**).

## Discussion

Through CFD simulations we observe two orders of magnitude difference in WSS between the lowest physiologically relevant ostium positions and the highest, indicating that the hemodynamic shear loads on the mouse AoV cusps during end diastole can vary greatly depending on ostia position. This gave us a unique opportunity to assess the effect of coronary ostium position, or lack thereof, on calcification in a mouse model of calcific AVD. When the ostia are high enough, as is often the case with the mouse RCO, the shear patterns on the aortic cusp surface during end diastole of the RCC and the NCC are nearly identical. Yet there is no correlation between coronary ostium position and calcification in either the LCC, NCC or the RCC in this mouse model of AVD. Similar to human patients, asymmetric pathological remodeling, where the NCC presented higher calcification burden, was conserved. This indicates that there may be another underlying factor that differentiates the NCC from its ostia bearing counterparts that predisposes it to higher calcific burden. Collagen and elastin fibers are known to serve as nucleation points for calcification [45–47], but our data suggests that the LCC contains more elastin and collagen than the other two cusps. This does not follow the asymmetric onset of calcification in the mouse model. Overall, the diet significantly reduced leaflet thickness and elastin abundance but had no effect on collagen abundance yet we were unable to uncover a leaflet dependent response. Assessing ECM content throughout different timepoints of the diet treatment and disease progression is necessary to assess the effect of ECM remodeling on asymmetric onset of AVD in this model.

Though previous literature suggests that the main difference in shear stresses on the aortic valve cusps would occur during diastolic loading, it is possible that the presence of the ostia asymmetrically affects systolic hemodynamic loading conditions as blood flows back into the sinuses through ventricular contraction and the venturi effect of the blood flow through the aorta. Asymmetric hemodynamic and mechanical loading during systole could also be driving these differences. We take a quasistatic steady state approach and only assess shear during end diastolic loading.

The extent to which this mouse model recapitulates the pathological disease progression in humans is unknown particularly in the context of CKD. Though calcification burden trends closely followed what is observed in human patients (biological sex and leaflet dependent trends), our ECM abundance and leaflet thickness quantifications it suggests that the remodeling in this mouse model is different from the classic inflammation driven AoV remodeling that incudes fibrosis and calcification. However, it is unknown if this classic pathological progression is conserved in CKD associated AVD progression in humans.

Through spatial transcriptomic analysis, we note both sex dependent and leaflet dependent differences in gene expression. Interestingly, canonical cell type specific phenotypic identifiers were amongst the top 10 DEGs when comparing the cusps to one another. *DCT*, *PMEL*, and *TYRP1* are all canonical melanocyte specific phenotypic markers which were in the top 10 downregulated genes in the LCC compared to the RCC and NCC. *MYH11,* a canonical phenotypic marker for smooth muscle cells and general cellular contractility, was in the top 10 upregulated genes in the NCC compared to the other two cusps. Also of note, *PALMD*, a recently identified susceptible gene for calcific aortic stenosis through transcriptome-wide association studies, was a top 10 upregulated gene in the NCC [48].

These findings were largely consistent with bulk RNA sequencing analyses where we were able to assess differential expression from 7 biological replicates and 21 different cusps. Among the top 10 DEGs in the NCC compared to the other two cusps, 8 genes were associated with smooth muscle cell contractility. The other 2 genes (*NOV*, and *SOST*) are both previously reported to be involved in aortic valve remodeling [49, 50]. DEG profiles showed melanocyte specific pathways upregulated in the NCC and downregulated in the LCC, while a contractile vascular smooth muscle cell phenotype was increased in the NCC and decreased in the LCC and RCC. GO biological process pathway “ossification” (GO:0001503) was upregulated in the RCC and downregulated in the LCC. Because these analyses were done on healthy adult WT mice, it suggests that there are intrinsic biological phenotypic differences between the three cusps that could exacerbate or protect against pathological remodeling under similar biomechanical and biochemical conditions. Whether these differences could be due to unaccounted for differences in stimuli across the 3 cusps, particularly during systole, is yet to be determined.

Our transcriptomic analyses were confined to healthy valves, rather than extended into the CKD+HP disease model. By focusing on baseline cusp identity, we were able to reveal intrinsic differences in contractile, extracellular matrix, and melanocyte-related programs, but we cannot determine how these signatures are reprogrammed once disease drivers such as inflammation, oxidative stress, or pro-calcific signaling are engaged. In calcific valve disease, pathways including TGF-β, Notch, Wnt, and osteogenic regulators become highly activated, and these inputs may converge differently with the baseline NCC, LCC, or RCC-specific phenotypes we describe. Thus, it remains unclear whether the cusp-specific phenotypic profiles act as a true priming mechanism that biases susceptibility to calcification, or whether it is rapidly remodeled by disease-induced cues to induce an “activated” cellular phenotype. Longitudinal profiling of cusps in diseased models will be required to dissect how intrinsic programs intersect with pathological signaling cascades, and whether specific molecular pathways uncovered here (e.g., SMC contractile or melanogenesis) persist as causal drivers or are supplanted during disease progression The AoV cusps are disproportionately infiltrated by different developmental origins during valvulogenesis, the endothelium, cardiac neural crest, and second heart field. The cardiac neural crest in particular infiltrates the NCC differently than the LCC and RCC [51]. Cushion cellularization (E9.5-E12.5) occurs prior to ostia origination (E14.5) in the primordial valve [52]. Thus, it is possible that under similar loading conditions the distinct cellular populations that arise from this asymmetric infiltration uniquely respond to similar biomechanical stimuli. Our computational deconvolution of the bulk RNA-seq data to determine cell-fractions within each leaflet aimed at informing this hypothesis better. Using scRNA-seq references and a deep autoencoder framework for tissue-adaptive mixture analysis (TAPE) we were able to compute the theorized cell fractions of previously identified major cell types within murine valvular tissues (interstitial cells, endothelial cells, immune cells, and melanocytes). The proportion of these cells were not evenly distributed among the leaflets suggesting that the asymmetries in developmental lineage patterning may persist into mature cell type differences in adulthood.

The quality of reference scRNA-seq has a great impact on bulk RNA-seq computational deconvolution. Single cell sequencing of mouse aortic valve leaflets is very difficult and requires pooling of multiple samples to get enough cells for analysis. For example, the scRNA-seq reference used for this deconvolution, Hulin et.al. 2019, pooled together 10 mice with both aortic and mitral valve tissues included. Though this is the best publicly available dataset which focuses on mouse AVL scRNA sequencing, it did not include other well established cell types found in the AVLs such as neurons and glial cells [53]. Thus, the computational deconvolution would be better informed by tissue specific, more robust and sensitive methods able to capture these cell types and require less samples to be pooled.

These findings could serve as a springboard to determine novel mechanisms of disease progression, beyond the classical VIC/VEC paradigm, and potential therapeutic targets to combat pathological ECM remodeling and calcification burden in patients with AVD.

## Abbreviations

AoV: aortic valve
AVD: aortic valve disease
AWSS: average wall shear stress
CAVD: calcific aortic valve disease
CFD: computational fluid dynamics
CKD: chronic kidney disease
CLR: centered log-ratio
ECM: extracellular matrix
LCC: left coronary cusp
LCO: left coronary ostium
NCC: non coronary cusp
RCO: right coronary ostium
RCC: right coronary cusp
STJ: sinotubular junction
WSS: wall shear stress

